# Enhancing the accuracy of genome-scale metabolic models with kinetic information

**DOI:** 10.1101/2024.07.11.597182

**Authors:** Jorge Lázaro, Arin Wongprommoon, Jorge Júlvez, Stephen G. Oliver

## Abstract

Metabolic models can be used to analyze and predict cellular features such as growth, gene essentiality, and product formation. There are several metabolic models but two of the main types are the constraint-based models and the kinetic models. Constraint-based models usually account for a large subset of the metabolic reactions of the organism and, in addition to the reaction stoichiometry, these models can accommodate gene regulation and constant flux bounds of the reactions. Constraint-based models are mostly limited to the steady state and it is challenging to optimize competing objective functions. On the other hand, kinetic models contain detailed kinetic information of a relatively small subset of metabolic reactions; thus, they can only provide precise predictions of a reduced part of an organism’s metabolism. We propose an approach that combines these two types of modeling to enrich metabolic genome-scale constraint-based models by re-defining their flux bounds. We apply our approach to the constraint-based model of *E. coli*, both as a wild-type and when genetically modified to produce citramalate. We show that the enriched model has more realistic reaction flux boundaries. We also resolve a bifurcation of fluxes between growth and citramalate production present in the genetically modified model by fixing the growth rate to the value computed according to kinetic information, enabling us to predict the rate of citramalate production.

**IMPORTANCE:** The investigation addressed in this manuscript is crucial for biotechnology and metabolic engineering, as it enhances the predictive power of metabolic models, which are essential tools in these disciplines. Constraint-based metabolic models, while comprehensive, are limited by their steady-state assumption and difficulty in optimizing competing objectives, whereas kinetic models, though detailed, only cover a small subset of reactions. By integrating these two approaches, our novel methodology refines flux bounds in genome-scale models, leading to more accurate and realistic metabolic predictions. Key highlights include improved predictive accuracy through more realistic flux boundaries, application to both wild-type and genetically modified *E. coli* for citramalate production, successful resolution of the bifurcation between growth and product formation, and broad applicability to other organisms and metabolic engineering projects, paving the way for more efficient bioproduction processes.

## INTRODUCTION

### Kinetic and constraint-based models

Metabolic models are mathematical representations of the metabolic reactions of an organism. Such models have been used to analyze key features of the organism, e.g. gene essentiality [1] and growth, and to develop new biotechnological milestones [2, 3, 4, 5].

Two popular modeling approaches for metabolism are kinetic models [6] and constraint-based models [7, 8]. While both approaches specify the stoichiometry of metabolic reactions, kinetic models provide detailed reaction rates based on metabolite concentrations and reaction kinetics, whereas constraint-based models only specify flux bounds based on stoichiometry and mass conservation constraints. Kinetic and constraint-based models are analyzed in different ways: integrating differential equations in kinetic models produces precise deterministic time trajectories of both the concentration of metabolites and the reactions’ rates, while constraint-based models can be analyzed as linear programming problems to produce a potential set of optimal fluxes over the reactions.

Kinetic models are difficult to develop because modeling and validating the kinetics of a metabolic reaction requires a large amount of information [9]. Furthermore, the integration of large non-linear ODE systems might be computationally complex. Therefore, such models are relatively small and tend to focus on central carbon metabolism or a particular pathway [10]. On the other hand, constraint-based models only specify the stoichiometry and the flux bounds of reactions. Given the simplicity of the constraints that they incorporate, i.e. just reaction stoichiometries, it is relatively easy to build large constraint-based models; consequently, genome-scale constraint-based models exist for several organisms [11, 12]. A disadvantage of these models is their lack of precision [13, 14, 15] and although genome-scale metabolic models have proven useful to predict specific phenotypes (e.g. growth rates in different conditions) the constraints that they usually include are not enough to accurately predict intracellular fluxes [16]. Moreover, estimating the values of competing objective functions, such as biomass and product formation, is not straightforward in constraint-based models.

Previous studies have attempted to bridge the gap between kinetic and constraint-based models. In [17], the authors examined the production of docosahexaenoic acid from the dinoflagellate *Crypthecodinium cohnii* by evaluating whether fluxes from independently simulated kinetic and constraint-based models were in agreement. However, this study does not integrate information from kinetic model simulations to enhance the constraint-based model, or vice versa. The work in [18] reviews methods to combine kinetic and constraint-based models and provides a toy example based on first principles. However, the toy example only has six reactions, substantially less than the number of reactions in recent kinetic and constraint-based models, therefore, applying their method to such models will be computationally expensive due to the exponential increase derived from different combinations of parameter values. The study further proposes narrowing the solution space, but this does not address pathway branching that can arise when there are competing objective functions in a constraint-based model. Both studies lack validation using metrics related to internal reaction fluxes. Thus, there is space for development of a computationally inexpensive way to use information from kinetic models to restrict the behavior of constraint-based models to make the output more realistic. In [19], the modeling formalism of Flexible Nets was employed to combine the dynamics of a bioreactor (macroscopic model) with a small metabolic network (microscopic model). In contrast to that approach, the present work simulates an independent kinetic model and uses the resulting fluxes to constrain reactions in a genome-scale model. This approach requires mapping reactions between two distinct models: the kinetic model and the constraint-based model.

### Citramalate: An industrially relevant metabolite

Citramalate, also known as 2-hydroxy-2-methylbutanedioate or hydroxypyruvic acid, is a dicarboxylic acid with a methylidene group in the middle of the carbon chain. Citramalate is not commonly encountered, but it is present in some bacteria, such as *Methanocaldococcus jannaschii* and fungi. It has been found out that it is an intermediate in certain metabolic pathways, including the biosynthesis of isoleucine in bacteria [20]. *M. jannaschii* expresses the gene *CimA3*.*7*, which codes for the enzyme citramalate synthase (EC:2.3.1.182) to produce citramalate. This enzyme catalyzes the reaction in which one molecule of acetyl-CoA, one molecule of pyruvate, and one molecule of water react to produce one molecule of (3R)-citramalate, one molecule of CoA, and a proton [21], see Equation (2).

Citramalate can be an alternative precursor to fossil fuel sources for the synthesis of industrially relevant compounds. Specifically, citramalate can be converted into methacrylic acid, a precursor for methyl methacrylate (MMA), which is in turn a component for the production of poly MMA (pMMA), a compound widely used in various industries including dentistry, electronics, and paints. Producing pMMA from citramalate thus results in a more sustainable and environmentally friendly process because the current commercial process for producing MMA involves using petroleum and natural gas. Extraction of such compounds is damaging for ecosystems, resulting in greenhouse gas emissions and hazardous waste generation [22].

To produce citramalate at an industrial scale, genetically modified organisms in bioreactors can be used. Although different microorganisms have been genetically modified to produce citramalate, *E. coli* is a good candidate for a citramalate factory because it produces biomass rapidly and can easily be genetically modified. In continuous and growth-limiting feed conditions, the bacteria produce 0.48 grams of citramalate per gram of glucose [23]. Therefore, mathematical modeling of citramalate production in genetically modified *E. coli* may lead to optimization of bioreactors, replacing fossil fuel precursors of MMA.

### Aims and objectives

We aim to bridge the gap between kinetic and constraint-based models. In particular, we show how to enhance a genome-scale constraint-based model with kinetic information obtained from a kinetic model in order to obtain a more realistic constraint-based model. We enriched the constraint-based model through the following steps: a) map reactions between the kinetic and the constraint-based models; b) compute fluxes of the kinetic model in steady-state simulations and; c) translate the bounds to the constraint-based model. To help map and translate flux bounds, we represent reactions graphically as Petri nets [24]. To assess the enhanced constraint-based model, Flux Balance Analysis (FBA) and Flux Variability Analysis (FVA)[25] were used.

We demonstrate our approach using a kinetic [26] and a constraint-based model [27] of *E. coli*. Concerning the FVA, we examined the number of reactions with increased or decreased variability (given by the flux variability value) after incorporating kinetic constraints into the original constraint-based model. We show that enhancing the constraint-based model with kinetic information results in the reallocation of reaction fluxes due to the incorporation of tighter constraints, as well as the activation of reactions that initially had no flux. Additionally, we extended these models to simulate the production of citramalate. An undesirable feature of the extended constraint-based model is a bifurcation which allows the model to fully direct the nutrient flux either towards the production of biomass or towards the production of citramalate. Thus, maximizing growth rate leads to null citramalate production, and similarly, maximizing citramalate production results in an unrealistic zero growth. We show that enriching this constrained-based model with kinetic information and fixing the growth rate solves the split of pathways problem, resulting in a more realistic value of citramalate production efficiency.

## MATERIALS AND METHODS

### Kinetic model

We use the *kinetic model* of *E. coli* introduced in [26]. This model captures the main central carbon pathways and accounts for 68 reactions and 77 metabolites which are located in 3 compartments: environment, periplasm and cytoplasm. The model represents glucose-limited conditions and is expressed as a set of ordinary differential equations (see [26] for details on the development of the model). Out of the 68 reactions, 48 include *V*_*max*_, the maximum flux that a reaction can carry under specific conditions, as a parameter. Of these reactions, 41 correspond to enzyme-catalyzed reactions (the remaining 7 reactions with an assigned *V*_*max*_ are either exchange reactions, the growth reaction, or the ATP_MAINTENANCE reaction, and hence, cannot be attributed to a particular enzyme). The *V*_*max*_ parameter can be perturbed to account for different cellular conditions arising from different enzyme concentrations, as expressed by Equation (1):

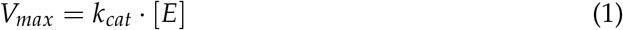

where [*E*] is the enzyme concentration and *k*_*cat*_ is the turnover number.

The kinetic model was analyzed by integrating its set of differential equations until the steady state was reached, evidenced by infinitesimal changes in substrate concentrations, usually after a simulation time of 3600 seconds. For this purpose, we used libRoadRunner 2.7.0 which is a C/C++ library that supports simulation of SBML based models [28].

### Constraint-based model

A *constraint-based model* is a tuple {ℛ, ℳ, 𝒮, *L, U*} where ℛ is the set of reactions, ℳ is the set of metabolites, 𝒮 ∈ ℝ ^|ℳ|×| ℛ|^ is the stoichiometric matrix, and *L, U* ∈ ℝ^|ℛ|^ are the lower and upper flux bounds, respectively, of the reactions.

The constraint-based model of *E. coli*, referred to as iML1515 and described in [8], was utilized in this study. The iML1515 model encompasses 2712 reactions, 1516 genes, and 1877 metabolites, which are located in three compartments, namely environment or extracelluar space, periplasm, and cytoplasm. Like any other constraint-based model, iML1515 is composed of a list of reactions, each of which specifies its reactants and products, along with constant lower and upper flux bounds.

### Petri nets to represent metabolic networks

Petri nets are a popular modeling formalism for dynamical systems [29, 30]. A Petri net is a directed bipartite graph with two types of vertices: *places*, which are depicted as circles, and *transitions*, which are depicted as rectangles.

Petri nets can be used to graphically represent metabolic networks [24]. In such a representation, metabolites are modeled by places and reactions are modeled by transitions. A metabolite that is a reactant in a reaction is connected with an arc from the metabolite to the reaction. Similarly, a metabolite that is a product in a reaction is connected with an arc from the reaction to the metabolite.

As an example, the Petri net in Figure **1** models the reaction for citramalate synthesis:

**FIG 1.**
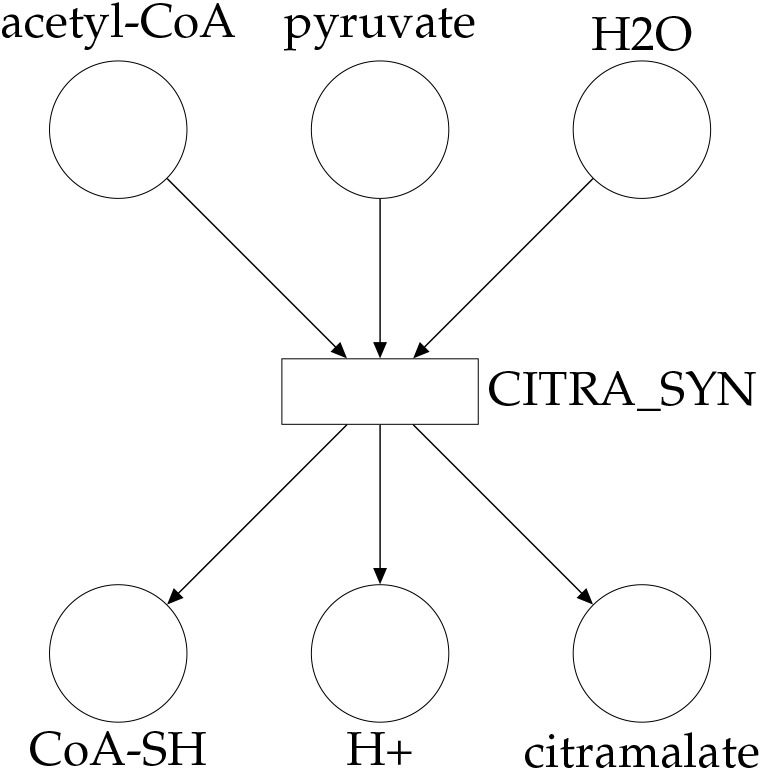
Petri net modeling the citramalate synthesis reaction specified in Equation (2). The reactants are in the upper part of the diagram and outgoing arcs come from these places, which means that there is a consumption of the reactants through the reaction *CITRA*_*SYN*. The products are those places with incoming arcs coming out of the transition.

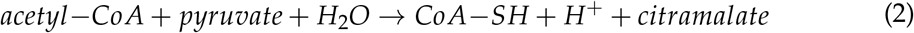

Metabolites *acetyl*−*CoA, pyruvate* and *H*_2_*O* are reactants so they are modeled as input places of transition *CITRA*_*SYN*, which models the reaction, and metabolites *CoA*−*SH, H*^+^ and *citramalate* are products, modeled as output places. A comprehensive Petri net is obtained once all the metabolites and reactions in the metabolic network are modeled with Petri net elements. Representing reactions as Petri nets helps us transfer flux bounds between constraint-based models and kinetic models.

### FBA and FVA simulations

Flux Balance Analysis (FBA) calculates steady-state metabolic fluxes in a network by optimizing an objective function (e.g., biomass production) subject to mass balance constraints. It assumes that the system is at a steady state, and the goal is to find a set of optimal fluxes that maximizes or minimizes a given objective function. FBA can be computed via the following linear programming problem:

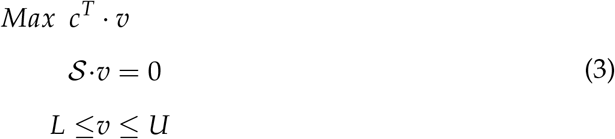

where *c* is a vector of weights indicating the contribution of each reaction to the objective function, 𝒮 is the stoichiometric matrix, *L* and *U* are lower and upper flux bounds, and *v* is the vector of fluxes of the reactions. The objective function, *c*^*T*^ · 𝓋, is maximized to find an optimal solution based on a set of optimal fluxes that satisfies the steady state condition, 𝒮 · 𝓋 = 0, and the flux bounds, *L* ≤ 𝓋 ≤ *U*.

Flux Balance Analysis (FBA) is most applicable under experimental conditions that facilitate a metabolic steady-state, such as nutrient-limited continuous cultures (chemostasts) where nutrient inputs (e.g., glucose, nitrogen) are tightly controlled, for instance, a chemostat in continuous culture where the growth rate is equal to the dilution rate at which nutrients are supplied. Controlled growth conditions (temperature, pH, oxygen) and data on nutrient consumption and byproduct secretion rates are essential for setting accurate constraints in the FBA model. Additionally, FBA models benefit from defining an objective function (e.g., maximizing growth or metabolite production) aligned with the specific experimental setup. These experimental conditions enable FBA to make reliable predictions about metabolic phenotypes under varying environmental and physiological scenarios.

Flux Variability Analysis (FVA) builds upon FBA by determining the range of feasible flux values for each reaction in the network. It calculates the maximum and minimum possible flux for each reaction while still satisfying the objective, whose optimized value is fixed, and mass balance constraints. FVA provides insight into the flexibility of metabolic pathways under different conditions, and can be computed by:

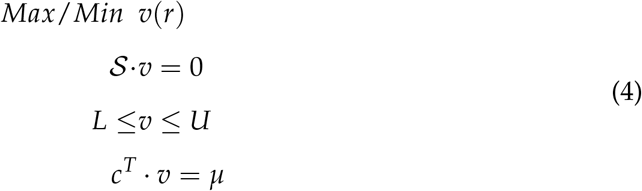

where *µ* is the solution of (3). If 𝓋 (*r*) in Equation (4) is maximized, then the maximum steady-state flux of reaction *r* is computed; similarly, if 𝓋 (*r*) is minimized, then the minimum steady-state flux of reaction *r* is computed.

FBA and FVA were used to elucidate the behavior of the enhanced constraint-based model. For the simulations, we used the value of glucose uptake in the kinetic model (0.23 *mMs*^− 1^) as the default constraint for the constraint-based model. The biomass reaction, *BIOMASS_Ec_iML1515_WT_75p37M*, was set as the objective function. COBRApy [31], a Python package to reconstruct constraint-based models, was used to translate the kinetic bounds and to run the FBA and FVA analyses.

## RESULTS

### Translating fluxes between the kinetic and constraint-based models

#### Translation of fluxes

To translate fluxes from the kinetic to the constraint-based model, for each reaction *r* in the constraint-based model, we define the flux bounds [*L* (*r*), *U* (*r*)] as follows:

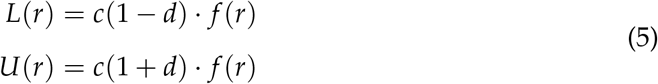

where:

- *f* (*r*) is the flux of reaction *r* obtained from the simulation of the kinetic model. This simulation was performed until the steady state was reached. Note that the all the computed fluxes of the kinetic model are positive (Table **2**).
- *L* (*r*) is the lower bound of reaction *r*.
- *U* (*r*) is the upper flux bound of reaction *r*.
- *d >* 0 is a real parameter that accounts for the degree of uncertainty of the model.
- *c* is a unit conversion factor.

The flux from the kinetic model, *f* (*r*), serves as the central estimate for the flux in the constraint-based model; *d* represents the relative uncertainty of the kinetic flux; and *c* is a unit conversion factor. For example, if *d* = 0.1 (i.e. uncertainty of 10%), then the lower bound of reaction r in the constraint-based model will be 0.9 times the flux of the reaction in the kinetic model, and the upper bound of the reaction in the constraint-based model will be 1.1 times the flux of the reaction in the kinetic model. Accounting for intervals of fluxes can be biologically interpreted as simulating perturbations in parameters that influence flux, such as enzyme abundance and the reaction’s catalytic constant (Equation 1).

In general, the kinetic bounds *L*(*r*) and *U*(*r*) obtained by applying Equation (5) were more restrictive than the original default bounds of the genome-scale model iML1515.

The unit conversion factor *c* exists because the kinetic and the constraint-based models use different units. Fluxes are in *mM s*^−1^ in the kinetic model [26], and in 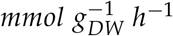 in the constraint-based model. In order to perform this unit conversion, we used the cell volume of 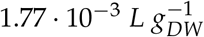 as specified in the kinetic model. In particular, a flux of *x mM s*^−1^ in the kinetic model is equivalent to the following quantity in 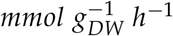

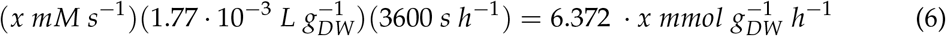

i.e., the conversion factor *c* is 6.372.

#### Mapping of reactions

As part of translating fluxes from the kinetic model to the constraint-based model, reactions in the kinetic model were mapped to their equivalents in the constraint-based model (Additional File 1). Even if the metabolic network of the constraint-based model is larger than that of the kinetic model, there might not be a one-to-one mapping of the set of reactions of the kinetic model onto the set of reactions of the constraint-based model. Some reactions of the kinetic model are identical to those of the constraint-based model, i.e., the sets of reactants, products and the stoichiometry are the same. In such a case, it suffices to convert the flux units, see (6), to translate the flux bounds of the kinetic model to the constrained-based model.

Non-identical reactions cannot be translated in the same way. Such reactions were classified in the following five categories, which were used to define equations that translate flux bounds to the constraint-based model (see Table **1** for an example reaction of each category):

1. **Difference in a small chemical species**: Reactions that differ in the species *H*^+^ or *H*_2_*O*.
2. **Acid hydrolysis**: Reactions in which the only difference is that *CO*_2_ is used in the constraint-based model, instead of 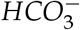 which is used in the kinetic model.
3. **Different reversibility**: Reactions that are reversible in one model and irreversible in the other model.
4. **Reversed**: Reactions that are reversed in the models, i.e. the metabolites that are reactants in one model are products in the other model.
5. **Subnetworks with different structure**: This category includes sets of reactions that perform the same chemical function but differ in the number of metabolites and in the connection pattern; thus, a one-to-one mapping is not possible.

For the reactions in categories *C*1, *C*2 and *C*3, the flux bounds of the kinetic model can be applied directly to the constraint-based model after unit conversion, i.e., as in Equation (5)

**TABLE 1.**
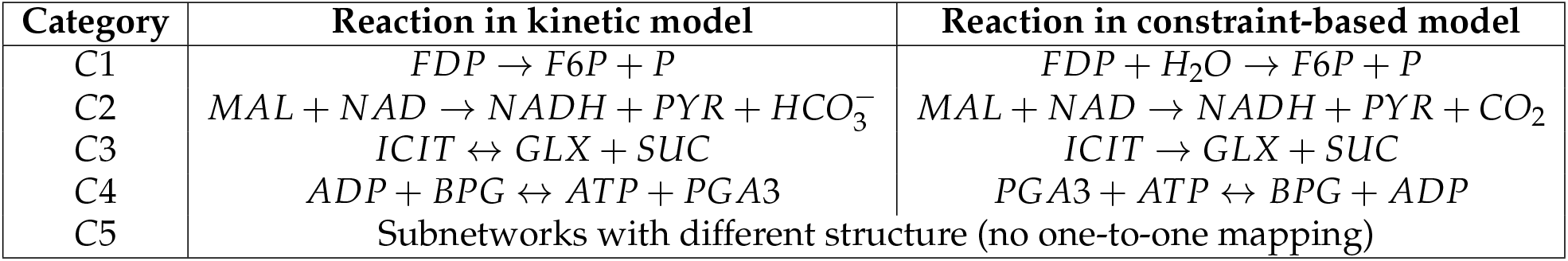
Categories of reactions to map fluxes from the kinetic to the constraint-based model.

| Category | Reaction in kinetic model | Reaction in constraint-based model |
| --- | --- | --- |
| C1 | $FDP \rightarrow F6P + P$ | $FDP + H_2O \rightarrow F6P + P$ |
| C2 | $MAL + NAD \rightarrow NADH + PYR + HCO_3^-$ | $MAL + NAD \rightarrow NADH + PYR + CO_2$ |
| C3 | $ICIT \leftrightarrow GLX + SUC$ | $ICIT \rightarrow GLX + SUC$ |
| C4 | $ADP + BPG \leftrightarrow ATP + PGA3$ | $PGA3 + ATP \leftrightarrow BPG + ADP$ |
| C5 | Subnetworks with different structure (no one-to-one mapping) |  |

In category *C*4, reactants and products are swapped, and hence, flux bounds of the constraint-based model must be computed as:

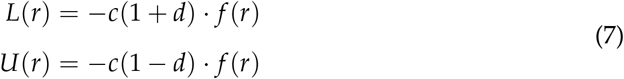

A procedure to compute flux bounds for reactions in category *C*5 is introduced in the Supplemental Text section (Supplemental Material).

### Enriching the constraint-based model predicts more realistic flux properties

To assess how changes in particular reactions can affect the set of optimal fluxes over the reactions of a genome-scale metabolic model, we sequentially restricted the flux bounds of the constraint-based model. Specifically, we sequentially added the bounds of 29 reactions following the glycolysis pathway (Table **2** in Supplemental Material), following Equation 5. We chose these reactions because we use a glucose-limited metabolic model and because adding constraints to more reactions causes the model to become infeasible as it overly restricts the linear programming solution space for FBA. After each reaction was added, we performed FBA and FVA, setting the parameter for the fraction of optimum as 0.999 for FVA. Setting this parameter in such a way is important because constraining the solution to the edge of the feasible state might lead to a reduced solution space which may result in a failure during the optimization process [32]. This approach allowed us to track specific changes in the behavior of the model as we implemented the kinetic bounds of each reaction.

To assess how the linear programming problem gets restricted and how the fluxes of the reactions distribute through the metabolic network, once the flux bounds of each reaction in Table **2** were added, we computed a) the FBA solution, b) the number of dormant reactions, and c) the feasible range *FV*(*r*) of fluxes of reactions before and after enriching the constraint-based model with the kinetic bounds. More precisely, *FV*(*r*) represents the potential flux variability of reaction *r* and is defined as:

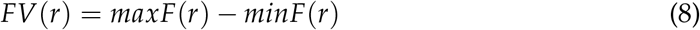

where *minF*(*r*) and *maxF*(*r*) are the minimum and maximum fluxes of reaction *r* computed by FVA.

A reaction *r* is said to be dormant when its steady-state flux is necessarily zero, i.e.:

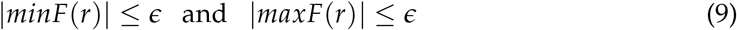

where *ϵ* is a small real quantity that accounts for the solver accuracy. To avoid numerical issues produced by the solver accuracy, *ϵ* has been set to 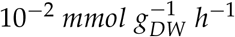 in our experiments.

#### Kinetic bound uncertainty of 3 %

To assess the effects of kinetic bound uncertainty values on the behavior of the model, we imposed uncertainty values *d* of 0.03, 0.06, 0.1 to capture a large range of model behaviors, ranging from the lowest level of uncertainty that results in a feasible solution to behaviors similar to the original model. To assess whether imposing a kinetic bound with uncertainty *d* = 0.03 makes the model more realistic, we computed the growth rate, the number of dormant reactions, and the variability of each reaction, as each bound was added to the constraint-based model. The uncertainty *d* = 0.03 represents the lowest level of uncertainty capable of mapping the 29 reactions listed in Table 2. A further reduction in uncertainty would lead to an infeasible solution prior to implementing these kinetic constraints. Figure **2** shows that as bounds were added, the growth rate was either maintained or decreased. This observation is expected because kinetic bounds involve a more constrained model.

**FIG 2.**
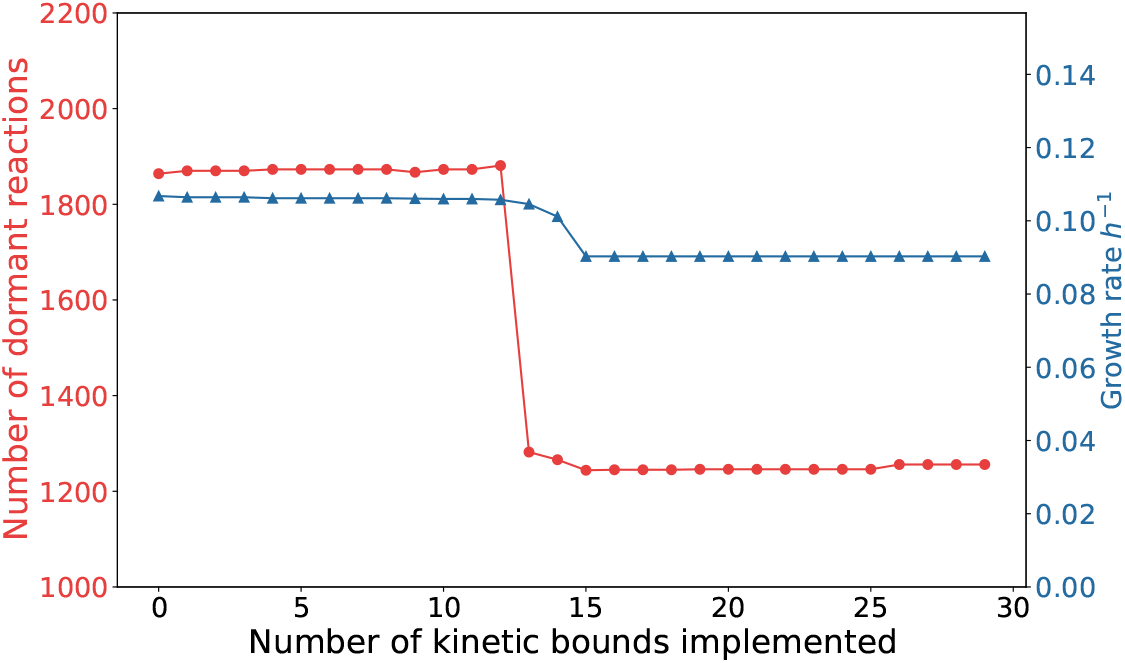
Growth rate (blue line) and number of dormant reactions (red line) with uncertainty parameter *d* = 0.03 as the number of implemented kinetic bounds increases.

**TABLE 2.** Reactions whose flux in the constraint-based model was limited to the kinetic bounds computed from the values in the second and third columns depending on the model studied. The kinetic flux values come from the steady state simulation of the kinetic model. Note that the names of the reactions correspond with the ones in the constraint-based model. Some of the names are different in the constraint-based model and the kinetic model, see Additional file 1 for a mapping of reactions between models.

| *ID* | *name* | *kinetic flux original model (mM s-1)* | *kinetic flux + citramalate (mM s-1)* |
| --- | --- | --- | --- |
| 1 | *PGI* | 0.168 | 0.170 |
| 2 | *PFK* | 0.184 | 0.197 |
| 3 | *FBA* | 0.184 | 0.197 |
| 4 | *TPI* | 0.184 | 0.197 |
| 5 | *GAPD* | 0.388 | 0.427 |
| 6 | *PGK* | 0.388 | 0.427 |
| 7 | *PGM* | 0.365 | 0.426 |
| 8 | *ENO* | 0.365 | 0.426 |
| 9 | *PYK* | 0.158 | 0.178 |
| 10 | *G6PDH2r* | 0.059 | 0.059 |
| 11 | *PGL* | 0.059 | 0.059 |
| 12 | *GND* | 0.042 | 0.040 |
| 13 | *RPE* | 0.017 | 0.027 |
| 14 | *RPI* | 0.025 | 0.014 |
| 15 | *TKT1* | 0.011 | 0.013 |
| 16 | *FBP* | 0.00012 | 0.00012 |
| 17 | *PPC* | 0.079 | 0.062 |
| 18 | *PPCK* | 0.111 | 0.044 |
| 19 | *PPS* | $1.5e - 5$ | $8.8e - 6$ |
| 20 | *ME1* | 0.022 | 0.032 |
| 21 | *PDH* | 0.383 | 0.257 |
| 22 | *CS* | 0.226 | 0.039 |
| 23 | *ACONTa* | 0.226 | 0.039 |
| 24 | *ACONTb* | 0.226 | 0.039 |
| 25 | *ICDHyr* | 0.128 | 0.024 |
| 26 | *AKGDH* | 0.111 | 0.024 |
| 27 | *SUCOAS* | 0.111 | 0.024 |
| 28 | *SUCDi* | 0.210 | 0.039 |
| 29 | *FUM* | 0.210 | 0.039 |

The number of dormant reactions is either maintained or decreased as the kinetic bounds are sequentially implemented. Small changes in the growth rate affect the number of dormant reactions. When reaction number 13 is added, the number of dormant reactions falls drastically, i.e. some reactions were dormant, but are awakened once the kinetic bounds are applied. This change is due to the combined effect of the implemented kinetic bounds and not to the addition of the bounds of a particular reaction. Including this combination of reaction 13’s kinetic bounds leads to a change in the FBA solution which was characterized by a reallocation of the optimal fluxes in order to find another optimal solution. In contrast, the original model without kinetic bounds is too loose and the optimization of the growth rate implies that most of the reactions are dormant or inactive, which is not realistic. Our simulations thus imply that the addition of kinetic bounds results in a more realistic set of optimal fluxes with fewer dormant reactions and a lower value of growth rate.

In order to assess the flux variability over reactions, *FV*(*r*) was used, see Equation (8). In particular, we computed the number of reactions whose *FV*(*r*) increased and the number of reactions whose *FV*(*r*) decreased after the addition of kinetic bounds with respect to the original constraint-based model. Figure **3** reports the number of reactions whose *FV*(*r*) increases and the number of reactions whose *FV*(*r*) decreases. Figure **3** is complementary to Figure **2** and shows that when the number of dormant reactions falls, several reactions increase their *FV*(*r*). This observation suggests that many of the dormant reactions increased their *FV*(*r*), so that metabolic flux is available through these reactions.

**FIG 3.**
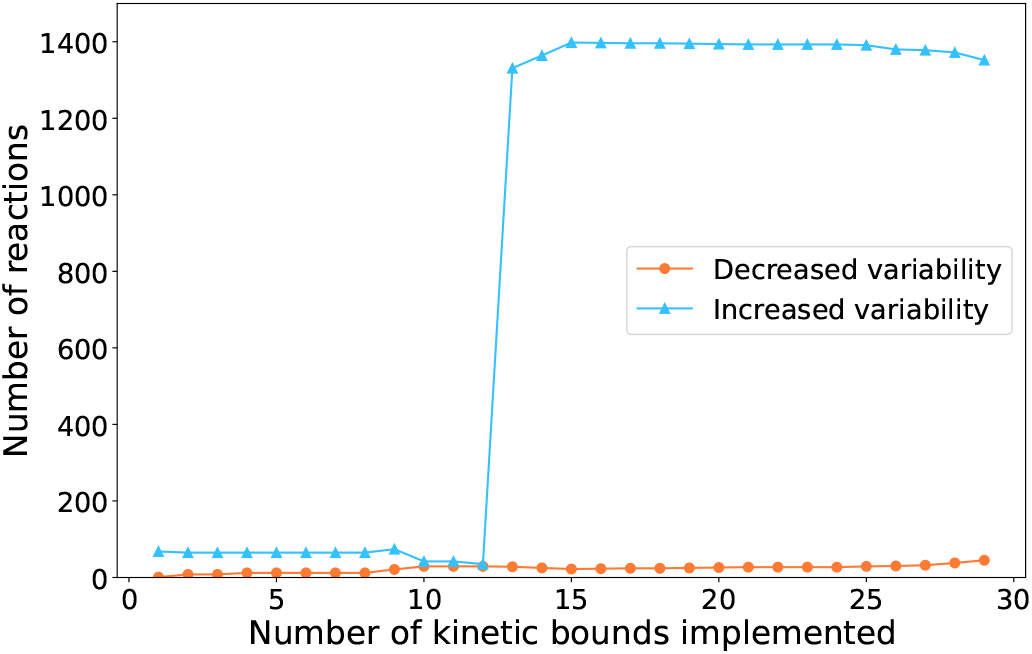
Number of reactions with more variability (blue line) and number of reactions with less variability (purple line) with respect to the original model as the number of implemented kinetic bounds increases with *d* = 0.03.

These results suggested that when the kinetic bounds of the reactions in Table **2** are added to the constraint-based model, the fluxes tend to reallocate in order to find a feasible solution. The limitations introduced by these new constraints do not allow flux values to achieve the same set of optimal fluxes as when the model has no flux bounds determined by the kinetic model. This leads to the following hypothesis: performing FBA on a model that has very loose constraints finds optimal solutions where many reactions in the model are dormant. Nevertheless, if we introduce additional constraints such as the kinetic bounds, the freedom of fluxes of certain reactions is reduced. Reactions that originally could get many different flux values, could no longer do so when they were constrained by the kinetic bounds. Thus, the original flux of a reaction has to reallocate through other reactions that were inactive. As a result, the optimal solution decreases and the optimal pathways that lead to the original solution are bounded, so new reactions have to activate and fluxes must be reallocated over the network to find a new optimal solution.

#### Kinetic bound uncertainty of 6 %

With an uncertainty of *d* = 0.06, the number of dormant reactions fell and the growth rate changed substantially when the kinetic bounds of reaction number 15 were added to the constraint-based model, see Figure **4**. Figure **5** shows that the change in the set of optimal fluxes occurs when the bounds of the reaction number 15 are implemented (the number of reactions that increase their *FV*(*r*) rises radically).

**FIG 4.**
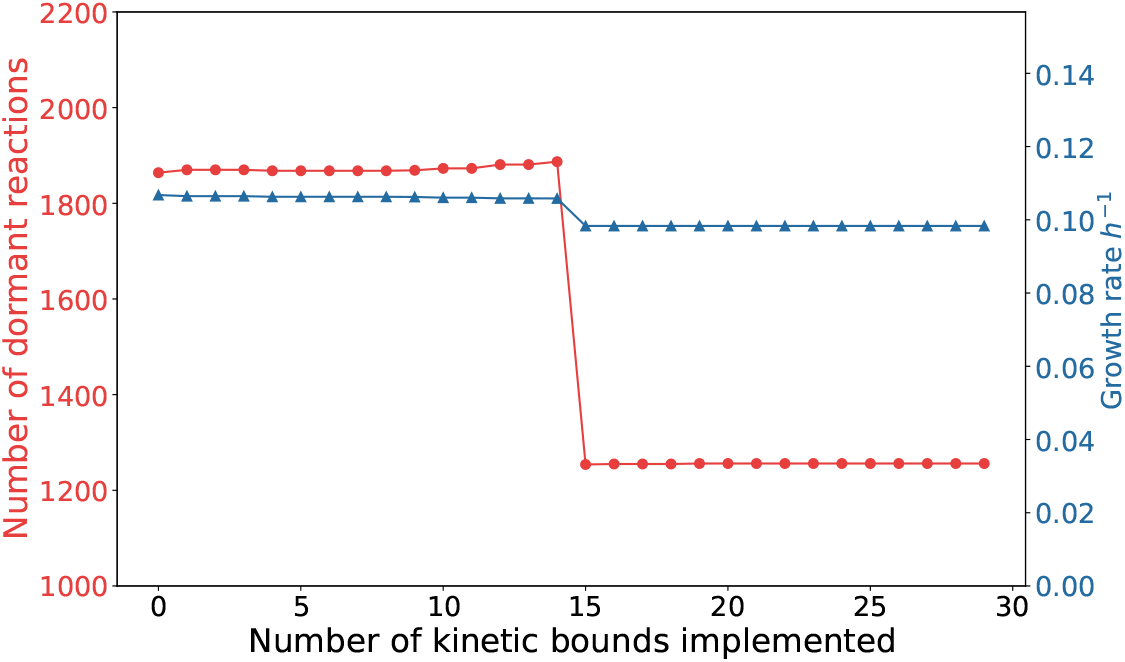
Dormant reactions (red line) and growth rate (blue line) as the number of implemented kinetic bounds increases when *d* = 0.06.

**FIG 5.**
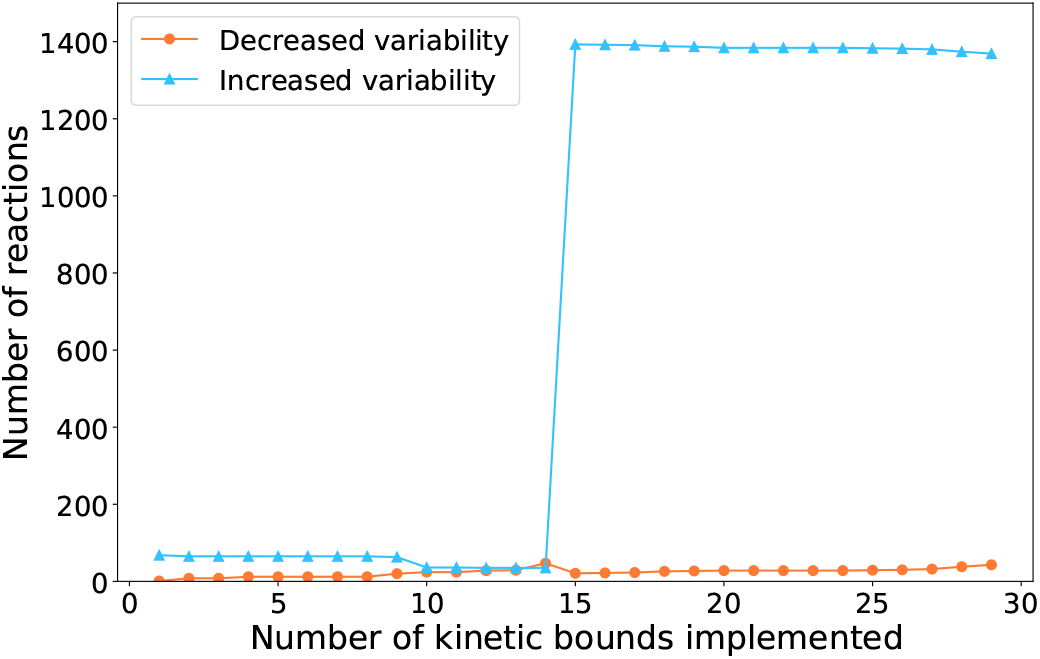
Number of reactions with more variability (blue line) and number of reactions with less variability (purple line) with respect to the original model as the number of implemented kinetic bounds increases when *d* = 0.06.

#### Kinetic bound uncertainty of 10 %

Finally, when the level of uncertainty was set to 10% (*d* = 0.1), neither the growth rate nor the dormant reactions are substantially affected, see Figure **6**. This phenomenon can also be seen in Figure **7**.

**FIG 6.**
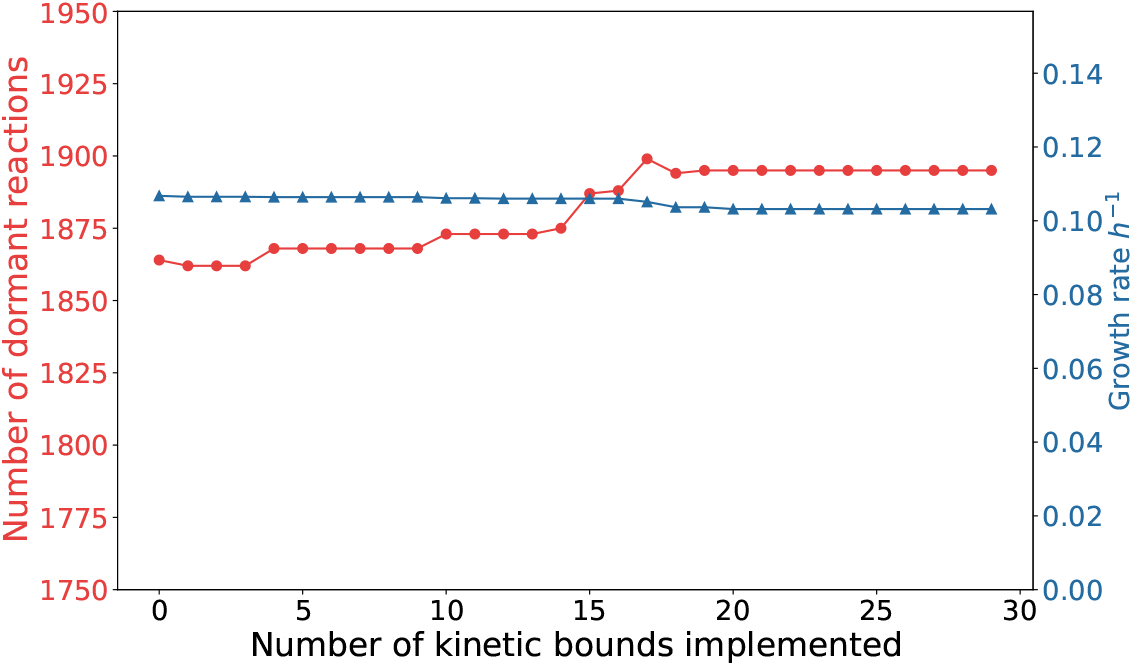
Dormant reactions (red line) and growth rate (blue line) as the number of implemented kinetic bounds increases when *d* = 0.1.

**FIG 7.**
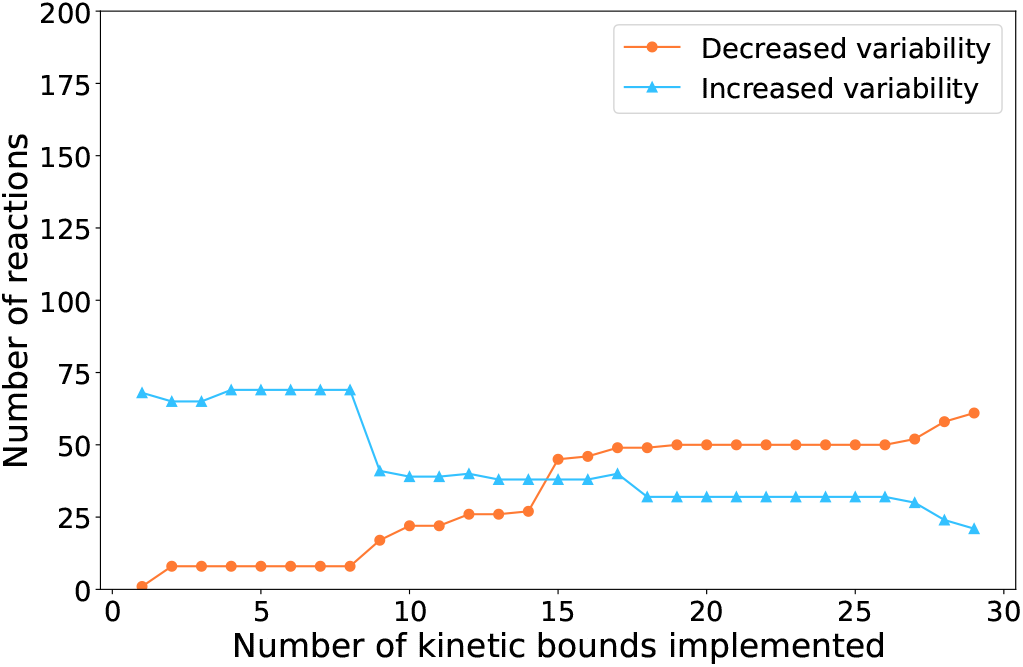
Number of reactions with more variability (blue line) and number of reactions with less variability (purple line) with respect to the original model as the number of implemented kinetic bounds increases when *d* = 0.1.

In general, when uncertainty is low, the number of dormant reactions falls after the addition of fewer kinetic bounds, leading to an earlier rise in reactions with increased variability. This is expected because a lower uncertainty in the kinetic bounds means that the model is more constrained, so the reallocation of metabolic fluxes happens earlier. This also results in reaching a lower optimal solution, i.e, growth rate, for the linear programming problem. In addition, giving more uncertainty to the kinetic bounds implies that we can implement the same or more kinetic bounds than for a lower level of uncertainty. This also makes sense because translating looser kinetic bounds entails that the model is being less constrained.

Additionally, to examine the variability of the reaction bounds, we computed the distribution of *FV*(*r*) values across reactions. The cumulative distribution of the *FV*(*r*) in Figure **8** shows that more than 90% of the total reactions had an *FV*(*r*) value lower than 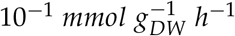 when FVA was performed on the original model and in the model enriched with kinetic bounds with an uncertainty of 10%. In contrast, less than 70% of reactions had an *FV* value lower than 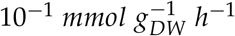 when FVA was performed on the kinetically constrained model under *d* = 0.06 and *d* = 0.03 uncertainty values. As *d* increases, the percentage of reactions for a given *FV* is higher. This means that higher levels of uncertainty in the translation of the kinetic bounds results in a lower *FV* and a small *d* is related to a higher *FV*. This agrees with the aforementioned finding: new reactions have to activate and fluxes must reallocate through the network to find a new optimal solution when kinetic constraints are added. Thus, a low uncertainty implies more reactions that increase their variability.

**FIG 8.**
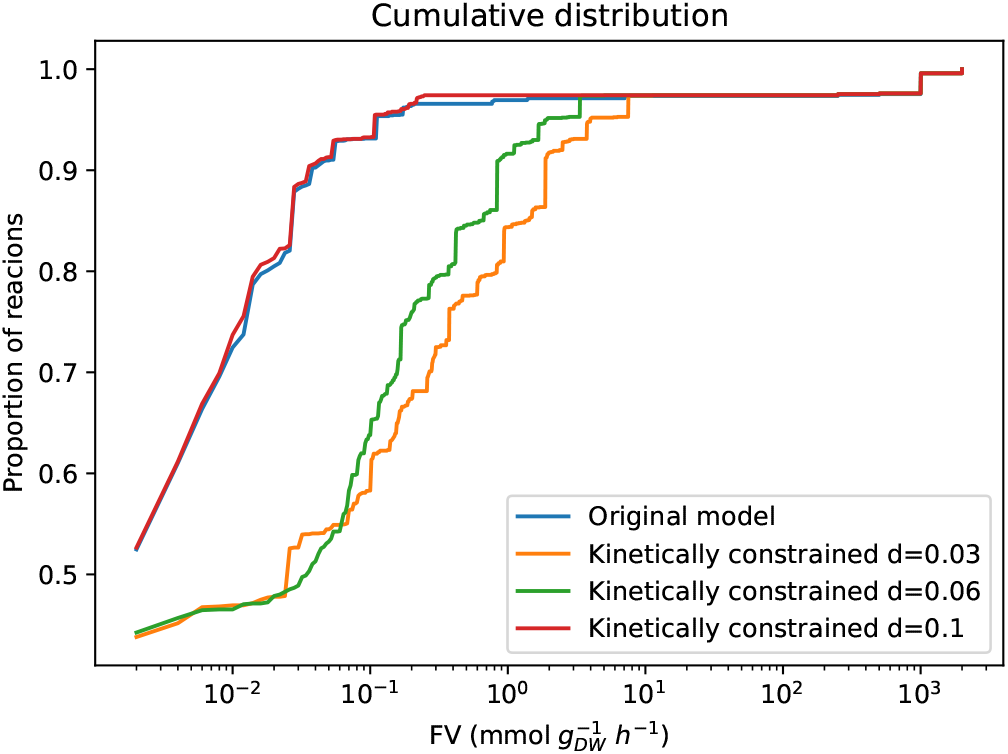
Cumulative distribution of *FV(r)* values in the original model (blue curve) and in the constraint-based model after the kinetic bounds for all reactions were added (kinetically constrained model) with *d* = 0.1 (red curve), *d* = 0.06 (green curve), and *d* = 0.03 (orange curve).

### Enriched constraint-based model to simulate citramalate production

#### Inclusion of citramalate production in the constraint-based model

One of the main motivations in this paper is to solve the problem that arises when there are two competing objective functions in the model. In a realistic model, there would be a trade-off between the growth rate and the production of citramalate. However, the carbon source is fully directed to the growth or to the citramalate production, depending on the objective function set to perform the FBA on the genome-scale model (Figure **9**). The kinetic model does not suffer from this problem since its differential equations impose positive fluxes in the reactions involved in the split of pathways between the growth and product formation as long as the concentration of the reactants is positive. Thus, to make the constraint-based model more realistic, we forced its growth rate to be in the interval defined by the growth rate generated by the kinetic model multiplied by the uncertainty, *d*. In other words, we added a new constraint specifying that the flux of the biomass reaction of iML1515 must be within the growth rate of the kinetic model plus or minus uncertainty *d*. This guarantees a strictly positive growth rate even if the objective function is the maximization of citramalate production.

**FIG 9.**
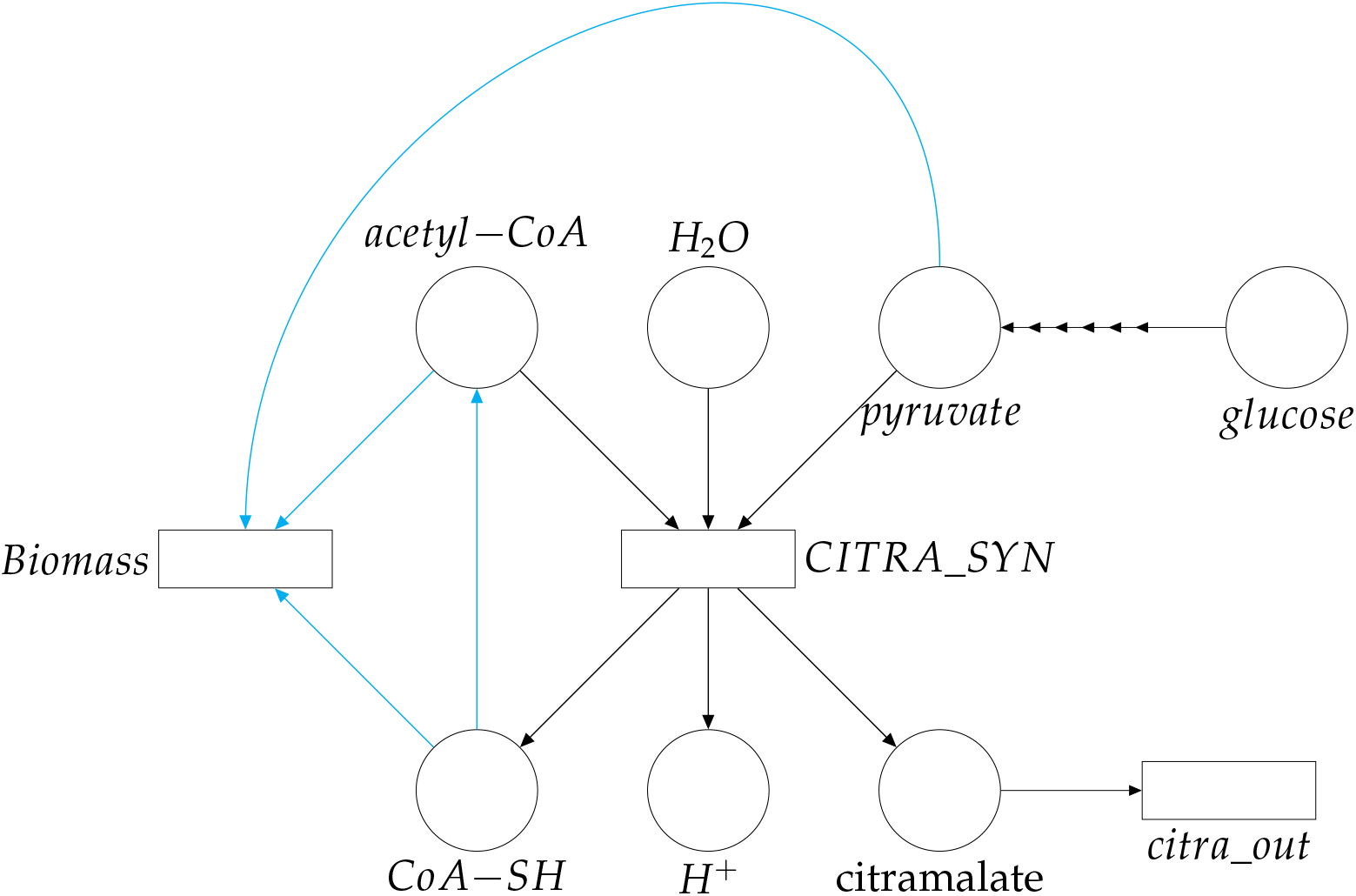
Simplified Petri net depicting location of bifurcation (cyan arrows) between growth and citramalate production. For simplicity, *citra_out* represents a subset of reactions (*CitraTransp1, CitraTransp2* and *EX_Citramalate*) added to the constraint-based model so that citramalate can leave the system. The arrowheads from glucose to pyruvate indicate the sequence of glycolytic reactions that transform glucose into pyruvate.

We modified the original constraint-based model of *E. coli* to model the synthesis and secretion of citramalate by the cell. The main reaction that was added to the model is (2) whose ID in the model is *CIMA*. Since the *E. coli* model has three compartments (cytosol, periplasm and extracellular), reactions that model the transport of citramalate from the cytosol to the periplasm (ID *CitraTransp1*) and from the periplasm to the exterior (ID *CitraTransp2*) were included as well. Finally, an exchange reaction (ID *EX_Citramalate*) that removes the citramalate from the extracellular compartment was added. Such an exchange reaction avoids the accumulation of extracellular citramalate and allows a steady state to be reached (Figure **9**). Subsequently, we chose this exchange reaction as the objective function in the FBA simulations.

#### Inclusion of citramalate production in the kinetic model

To model the synthesis of citramalate using the kinetic model, we added a new reaction, *CITRA_SYN*, to the model. The stoichiometry of this reaction is given by (2) and it follows the Michaelis-Menten kinetic law (10):

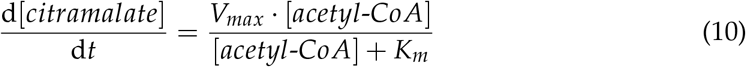

which assumes that pyruvate is saturating [33, 34, 35]. The *K*_*m*_ was set to 0.495 *mM* [36] and the *V*_*max*_ was calculated from the maximum activity of *CimA3*.*7* and the cell volume. More precisely, given that the maximum activity of *CimA3*.*7* in one cell is 24.34 · 10^−10^*nmol*/*s* [36], and that the volume of the cell is approximately 6 · 10^−16^ *L* [37], the *V*_*max*_ of *CimA3*.*7* can be calculated as:

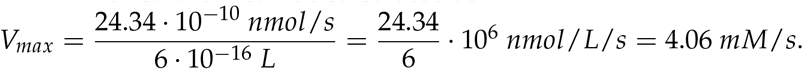

#### Effect of varying kinetic bounds in the genome-scale model

We enrich the constraint-based model with the citramalate synthesis reaction, with information from the equivalent kinetic model as we did previously for the wild-type constraint-based model. Figures **10**, **12** and **14** report the number of dormant reactions and the maximized citramalate production flux after implementing kinetic bounds in the wild-type model. They all show a pronounced fall in the number of dormant reactions, which coincides with the first substantial reduction in the citramalate production flux. Notice that the kinetic bounds have been implemented following the order of glucose metabolism, see Table **2**. Section “On the sequence of implemented kinetic bounds” in the Supplemental Material addresses the effect of a different implementation order on the results.

**FIG 10.**
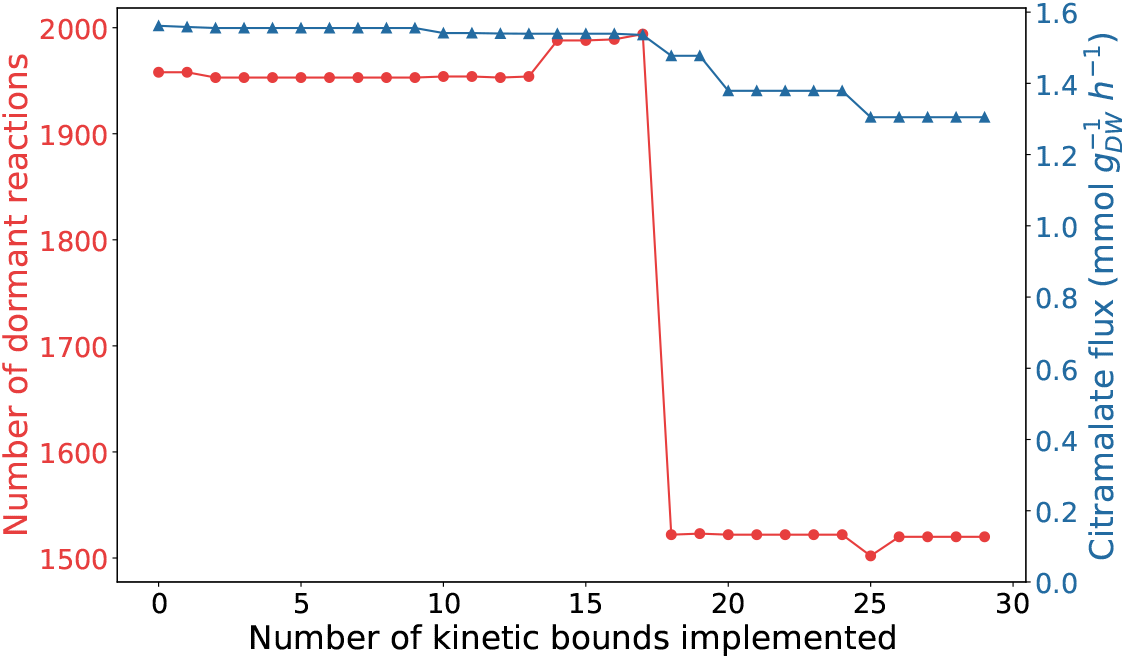
Dormant reactions (red line) and citramalate production flux (blue line) as the number of implemented kinetic bounds increases when *d* = 0.03.

**FIG 11.**
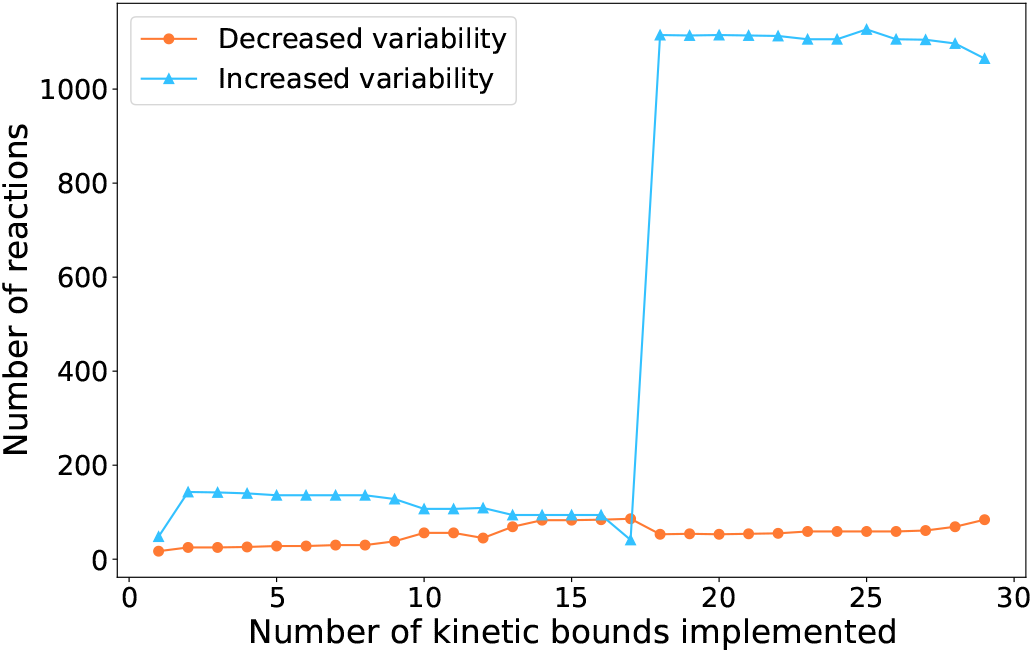
Number of reactions with more variability (blue line) and number of reactions with less variability (purple line) with respect to the original model as the number of implemented kinetic bounds increases when *d* = 0.03.

**FIG 12.**
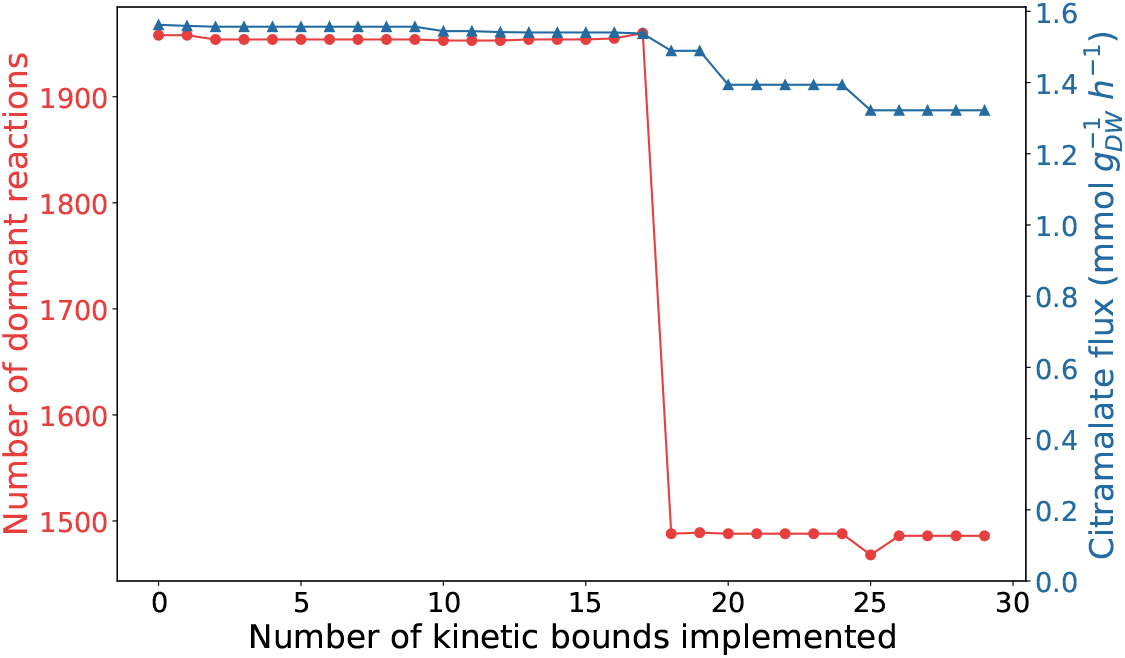
Dormant reactions (red line) and citramalate production flux (blue line) as the number of implemented kinetic bounds increases when *d* = 0.06.

Figures **11**, **13** and **15** show the number of reactions whose flux variability increases (blue curve) and decreases (red curve) after including the kinetic bounds with a level of uncertainty of 3%, 6% and 10% respectively. In all plots, adding kinetic bounds leads to more reactions that increase their flux variability in comparison to the flux variability that they exhibit when the solution is computed using the model without additional kinetic constraints.

**FIG 13.**
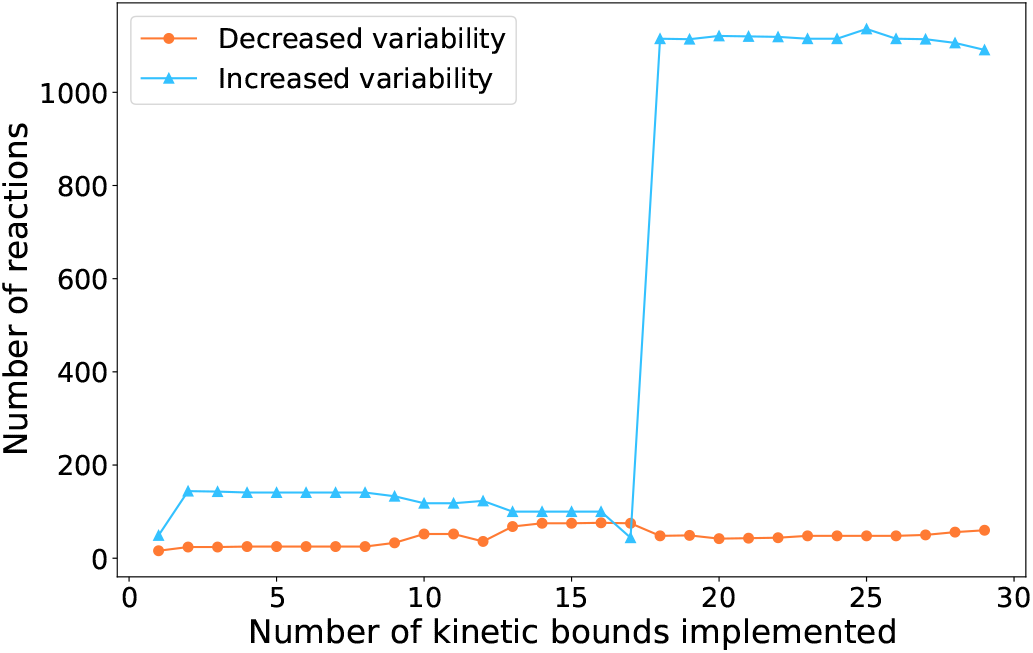
Number of reactions with more variability (blue line) and number of reactions with less variability (purple line) with respect to the original model as the number of implemented kinetic bounds increases when *d* = 0.06.

**FIG 14.**
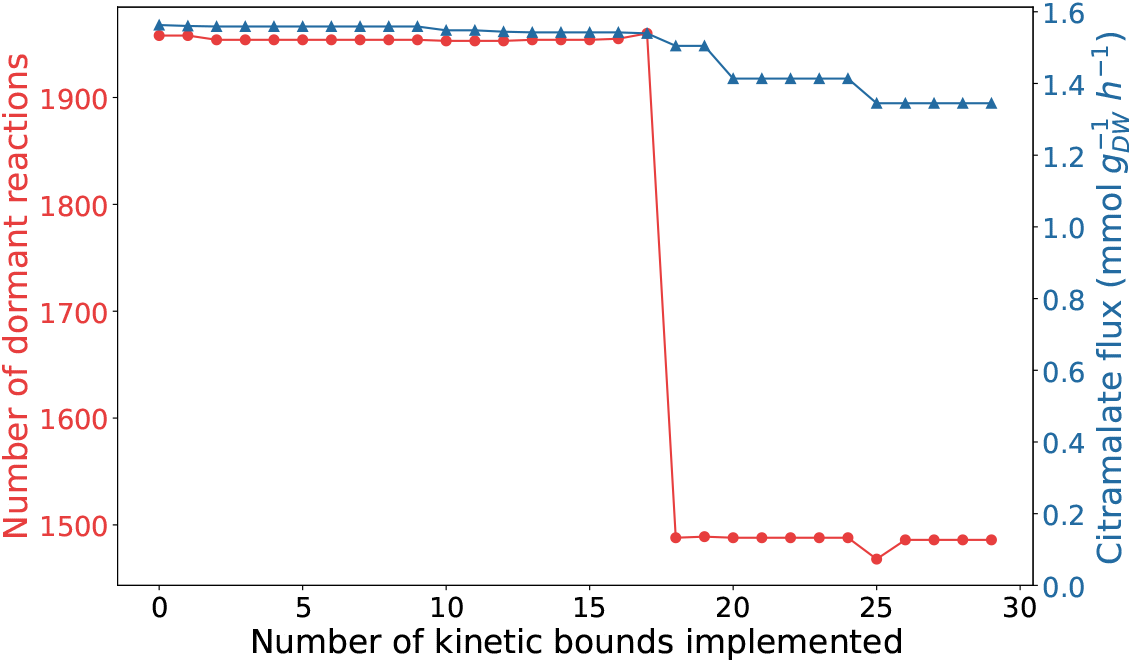
Dormant reactions (red line) and citramalate production flux (blue line) as the number of implemented kinetic bounds increases when *d* = 0.1.

**FIG 15.**
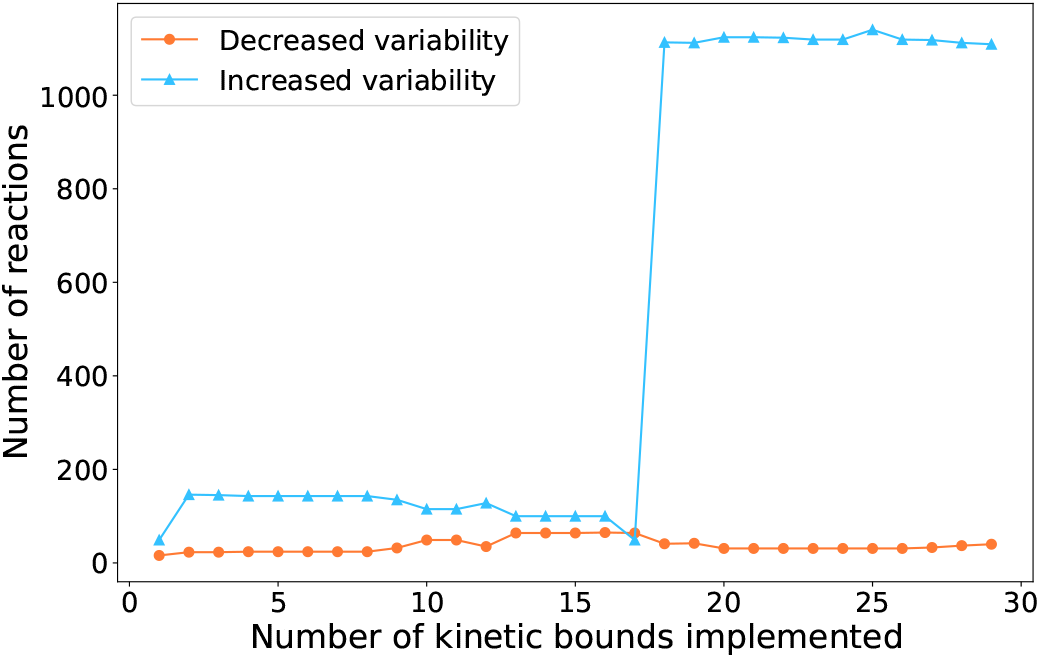
Number of reactions with more variability (blue line) and number of reactions with less variability (purple line) with respect to the original model as the number of implemented kinetic bounds increases when *d* = 0.1.

Once again, to assess the variability of the reaction bounds, we calculated the distribution of *FV*(*r*) values across reactions. Figure **16** shows that lower levels of uncertainty imply more reactions with higher variability, which is in agreement with Figure **8**. In summary, lower levels of uncertainty imply more reactions having higher variability.

**FIG 16.**
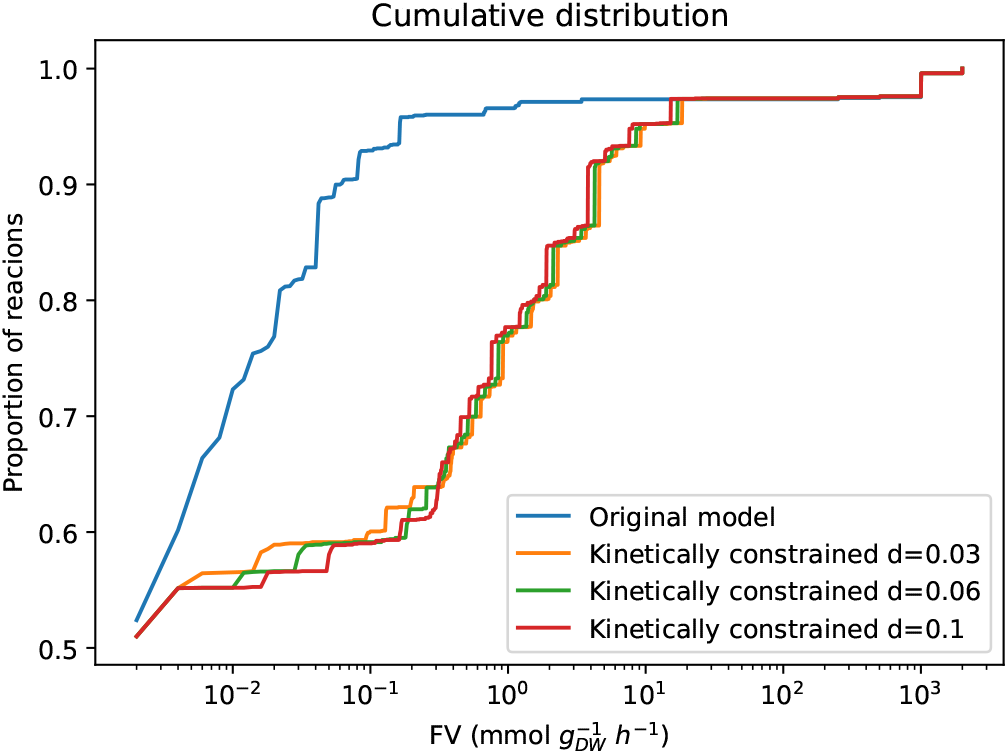
Cumulative distribution of *FV(r)* values in the enriched original model (blue curve) and in the constraint-based model after the kinetic bounds for all reactions were added (kinetically constrained model) with *d* = 0.1 (red curve), *d* = 0.2 (green curve), and *d* = 0.3 (orange curve).

#### Validation of citramalate efficiency in the kinetically constrained model

In order to validate the kinetically constrained model, we compared the predictions of the model with the experimental values reported in [23]. In our simulations the flux of citramalate, and hence the conversion efficiency, depends on the uncertainty used to implement the kinetic bounds in the model, see Figure **17**.

**FIG 17.**
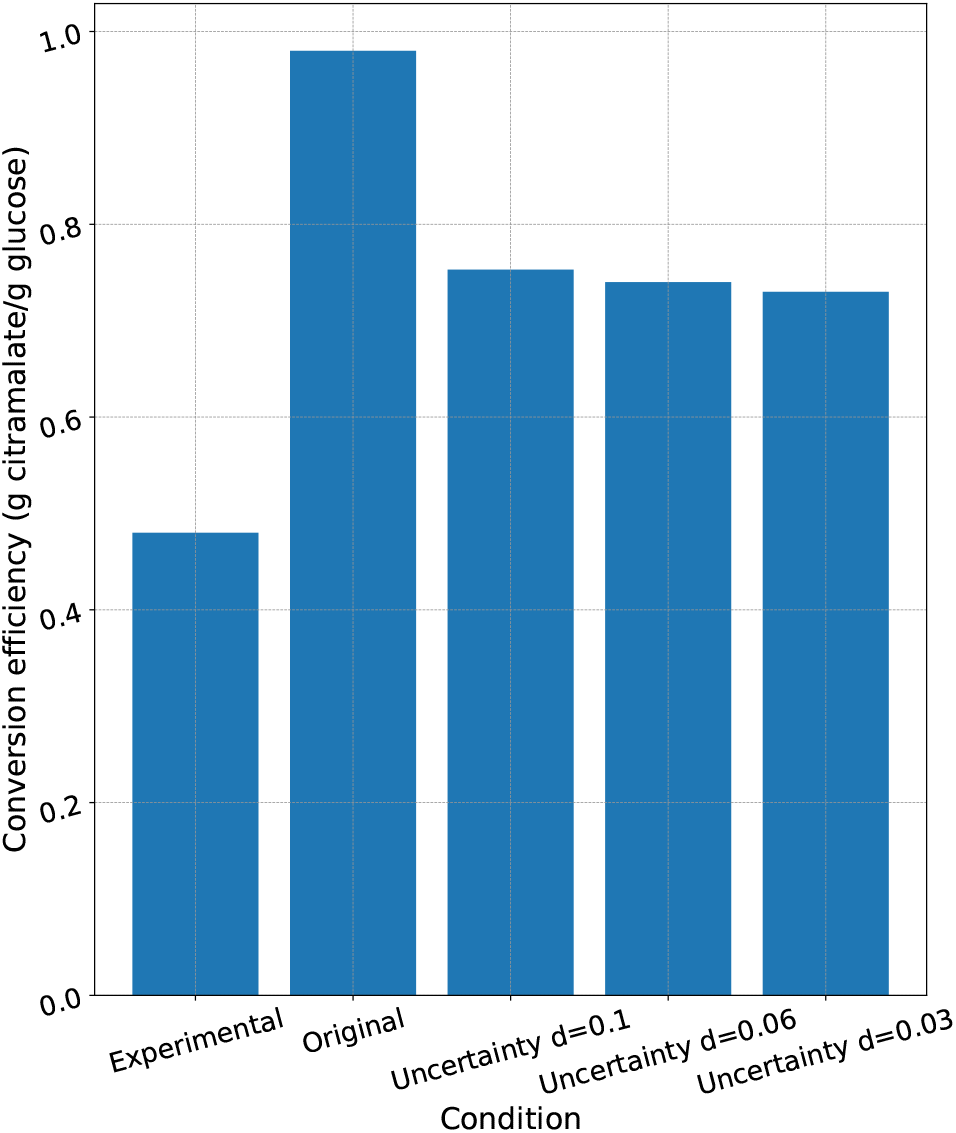
Conversion efficiencies from glucose to citramalate computed under different conditions where: *Experimental* corresponds to the experimental value for the conversion efficiency in *E. coli* according to [23] where the flow rate was manually adjusted as needed to sustain a pseudo-exponential growth rate of about 0.25 *h*^−1^ and to prevent excess glucose accumulation in the culture; *Original* is the conversion efficiency in the computational model iML1515 without additional constraints performing FBA; the rest of the conditions refer to the enriched computational model with kinetic bounds integrated with different levels of uncertainty (0.1, 0.06, and 0.03).

As expected, the glucose conversion into citramalate is highest for the original model, i.e. without kinetic bounds translated to the constraint-based model. This value likely overestimates the efficiency due to the lack of realistic constraints in the model. Thus, implementing the kinetic bounds into the constraint-based model decreased the value of the solution and brought the conversion efficiency closer to the experimental one. The efficiency progressively decreased as the level of uncertainty was reduced.

## DISCUSSION

Although constructing constraint-based models is straightforward, constraint-based models are imprecise and unreliable because they lack kinetic and proteomic information. Stoichiometry and flux bounds of reactions are the only constraints of constraint-based models. The default flux bounds assigned to most reactions are very loose and fail to realistically constrain the fluxes of reactions.

Therefore, we aim to make more realistic constraint-based genome-scale models by combining the properties of the former and kinetic models. Here we evaluate a genome-scale model of *E. coli* that we enrich with the data obtained from a simulation of the steady state of a kinetic model. The kinetic model is smaller than the constraint-based model and holds the kinetic information for the reactions involved in the carbon metabolism. We also create a new model that can produce citramalate based on kinetic parameters that were previously determined experimentally.

A key challenge in reconstructed networks is the occurrence of dormant reactions, which are constrained to zero flux and therefore remain inactive under steady-state conditions. The reduction of the number of non-functional reactions has been used to validate models as they provide better coverage of the space of metabolic reactions and exhibit fewer dormant reactions (and hence, considerably more reactions that can carry a flux different from zero) [38, 39]. Here, we show that the enriched model, with and without citramalate synthesis, is more realistic because adding new kinetic bounds to these models results in a decrease in the number of dormant reactions coupled to a generalized increase in the variability of the fluxes of the reactions. As kinetic constraints are added, the best ways to obtain the optimal solution are restricted and hence alternative, and probably less efficient, routes have to be activated in order to find a new optimal solution. This is very much more realistic than a non-constrained model where the objective function can be maximized in such a way that no flux is limited. Furthermore, we demonstrate that increased uncertainty of kinetic bounds leads to a model that behaves more closely to the unconstrained model. In contrast to other papers such as [16, 40], in some cases, adding kinetic constraints to a genome-scale model can increase the flexibility of reaction fluxes [41]. This effect arises when kinetic constraints unblock alternative pathways or allow previously restricted reactions to carry flux, effectively expanding the variability of the metabolic fluxes. Rather than strictly reducing variability, kinetic constraints can sometimes enable a redistribution of fluxes that activates additional metabolic routes under specific conditions.

In the extended citramalate-producing constraint-based model, there is a bifurcation that enables the model to exclusively channel nutrient flux either towards biomass or citramalate production. We resolved this split of pathways problem by fixing the growth rate to the value obtained from the kinetic model with the citramalate synthesis reaction added. Additionally, we show that implementing kinetic bounds with a level of uncertainty leads to the model predicting the efficiency of converting glucose to citramalate that is closer to experimental observations. It should be noted that, although constraining a constraint-based model with outputs from a kinetic model can improve model realism, this approach depends on the integrity and accuracy of the kinetic model itself, i.e. any modeling errors in the kinetic model will be carried over to the enriched constrained-based model. The citramalate production and growth rate trade-off is more realistic in the enhanced model, as it addresses the split of pathways that occurs in real life. Additionally, the presence of minimal dormant reactions aligns with realistic conditions, and the incorporation of kinetic constraints reflects resource allocation. Consequently, our enhanced model provides a more accurate prediction of growth and citramalate productivity in *E. coli* for industrial applications.

## Supporting information

https://github.com/jlazaroibanezz/citrabounds

## ACKNOWLEDGMENTS

The authors are grateful to Andrew Yakoumetti and Charlotte Green, University of Nottingham, UK, for their work to determine the values of the kinetic parameters in (10), and, especially, Andrew Yakoumetti for helpful discussions.

## DATA AVAILABILITY STATEMENT

The Python code generated for this work is available at: https://github.com/jlazaroibanezz/citrabounds

## SUPPLEMENTAL MATERIAL

### Supplemental Text

#### Subnetworks with different structure

Because one-to-one mapping is not possible for reactions in category *C*5, the flux bounds of the kinetic model cannot be applied directly to the constraint-based model.The three subnetworks in the kinetic model that belong to category *C*5 have different structures and, hence, will be considered separately. In the following graphical representations, a Petri net transition depicted as a double rectangle models a reversible reaction.

**Subnetwork 1**. The transformation of periplasmic glucose (*GLPc*) and phosphoenolpyruvate (*PEP*) into glucose-6-phosphate (*G6P*) and pyruvate (*PYR*) is modeled by 5 reactions, *PTS_0, PTS_1, PTS_2, PTS_3* and *PTS_4*, in the kinetic model, see Figure **18**, and by one reaction, *GLCptspp*, in the constraint-based model, see Figure **19**.

In the steady state, the overall incoming flux to a metabolite must be equal to its overall outgoing flux. Thus, the following equation will hold in the kinetic model:

**FIG 18.**
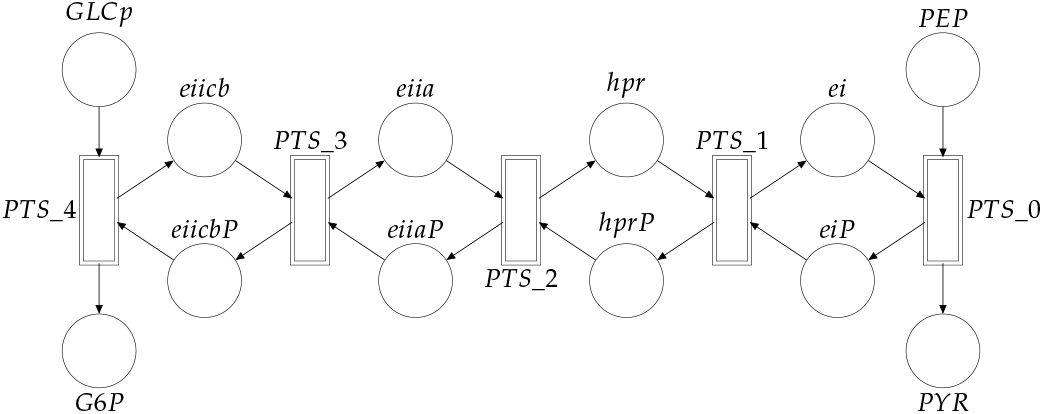
Petri net representation of the **Subnetwork 1** in the kinetic model, where the reactions are *PTS*_0, *PTS*_1, *PTS*_2, *PTS*_3 and *PTS*_4, the reactants are periplasmic glucose (*GLCp*) and phosphoenolpyruvate (*PEP*), and the products are glucose-6-phosphate (*G*6*P*) and pyruvate (*PYR*). The rest of the places represent intermediates participating in the reactions.

**FIG 19.**
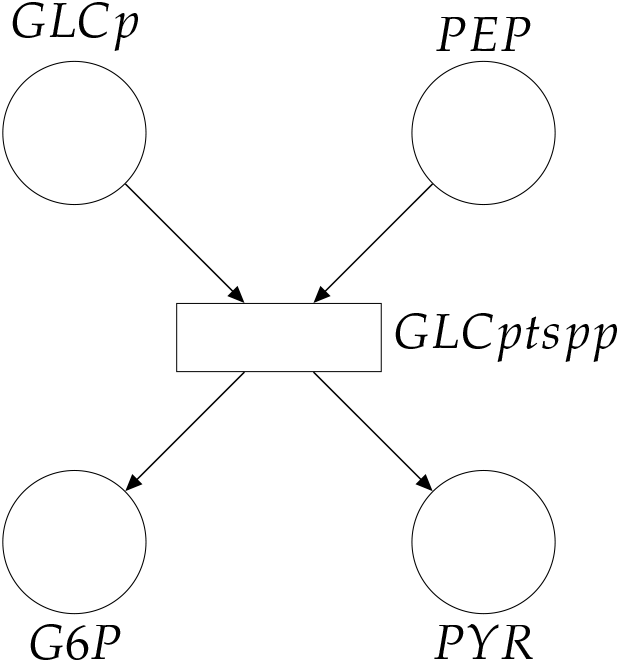
Petri net representation of the **Subnetwork 1** in the constraint-based model, where *GLCptspp* is the only reaction (notice that this reaction corresponds with the reactions in Figure **18**. of the kinetic model). The reactants of the reactions are periplasmic glucose (*GLCp*) and phosphoenolpyruvate (*PEP*), the products are glucose-6-phosphate (*G*6*P*) and pyruvate (*PYR*).

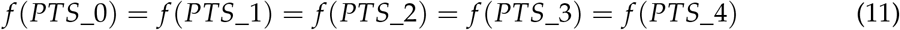

Hence, the flux of any reaction in Figure **18** can be taken to bound reaction *GLCptspp* in the constraint-based model. From Equation (5), appropriate lower and upper flux bounds for reaction *GLCptspp* can be computed by:

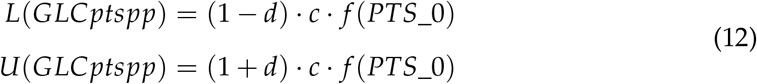

**Subnetwork 2**. The transformation of ubiquinone-8 (*Q*) and succinate (*SUC*) into fumarate (*FUM*) and ubiquinol-8 (*QH2*) is modeled by two reactions (*SDH* and *SQR*) in the kinetic model, see Figure **20**, and by one reaction (*SUCDi*) in the constraint-based model, see Figure **21**. Following the reasoning for *Subnetwork 1*, the two reactions of the kinetic model must have the same flux in the steady state. Thus, it holds:

**FIG 20.**
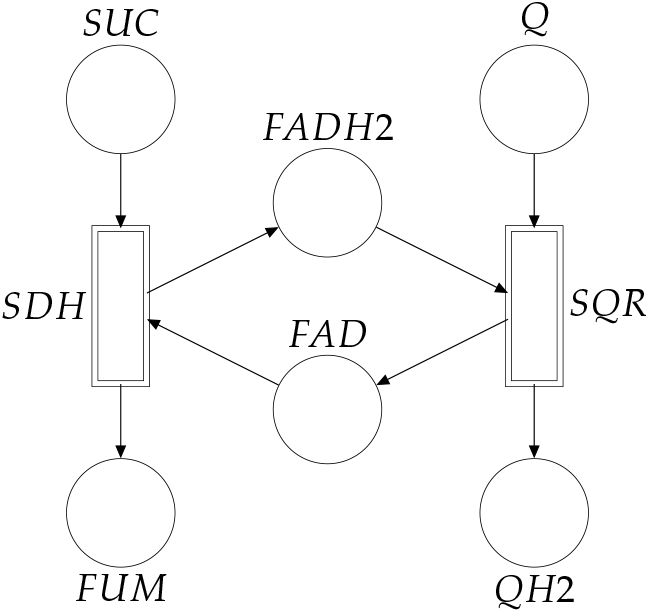
Petri net representation of the **Subnetwork 2** in the kinetic model.

**FIG 21.**
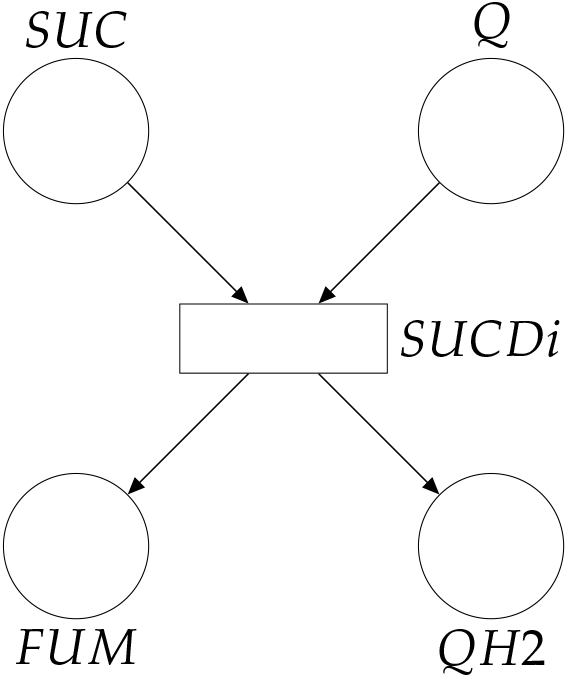
Petri net representation of the **Subnetwork 2** in the constraint-based model.

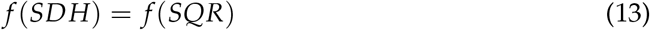

and the following bounds for the reaction *SUCDi* in the constraint-based model are obtained:

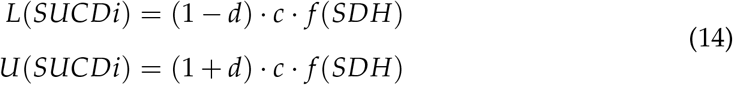

**Subnetwork 3**. The pentose phosphate pathway (PPP) is modeled differently in the kinetic and the constraint-based model. Namely, it is modeled by reactions *X5P_GAP_TKT, S7P_R5P_TKT, F6P_GAP_TAL, S7P_E4P_TAL* and *F6P_E4P_TKT* in the kinetic model, see Figure **22**; and by *TKT1, TKT2* and *TALA* in the constraint-based model, see Figure **23**. To map flux bounds in these subnetworks, we observe that various reactions in both models are equivalent, namely:

**FIG 22.**
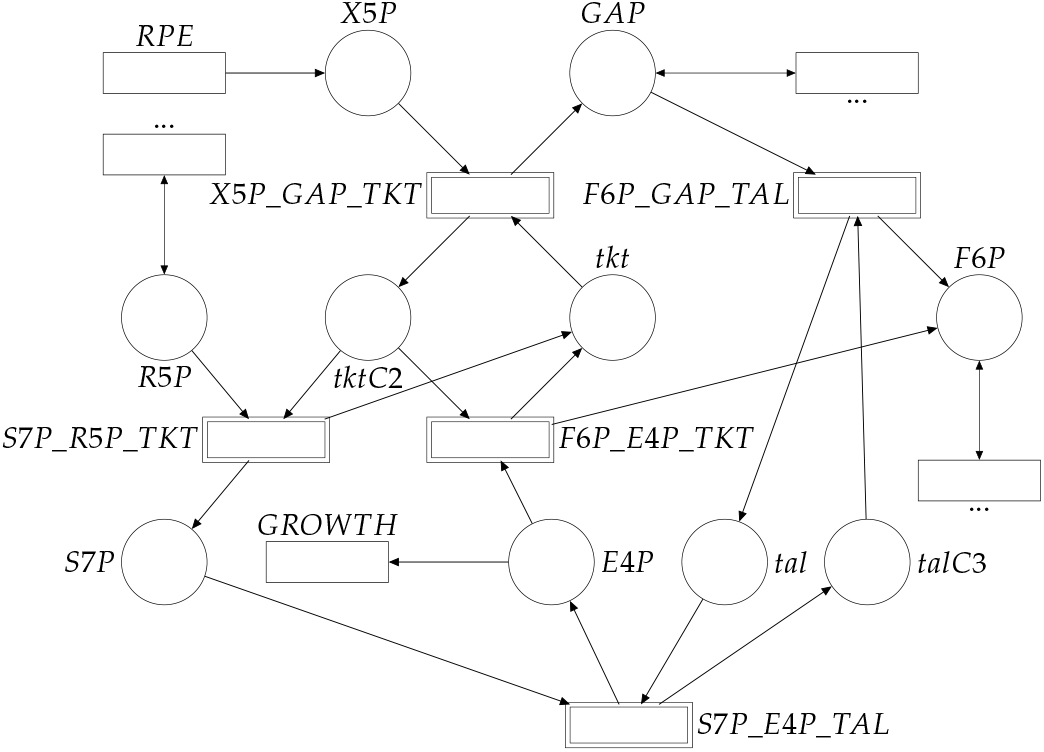
Petri net representation of the **Subnetwork 3** in the kinetic model. Reactions with a “…” as ID denote multiple reactions. If an arrowhead points to a place, one or multiple reactions produce the metabolite; if an arrowhead points to a transition, the metabolite is consumed by multiple reactions, and if the arrow is bidirectional one or multiple reactions consume or/and produce the metabolite.

**FIG 23.**
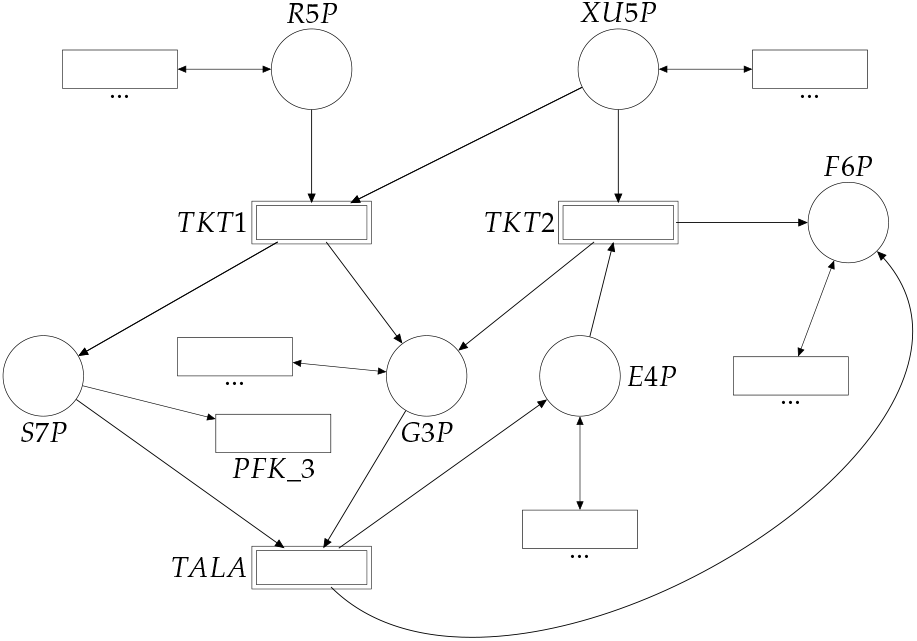
Petri net representation of the **Subnetwork 3** in the constraint-based model.

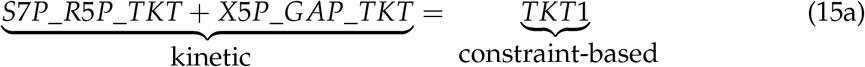

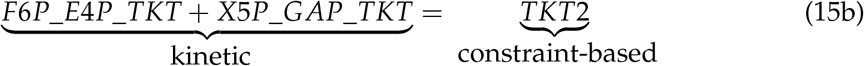

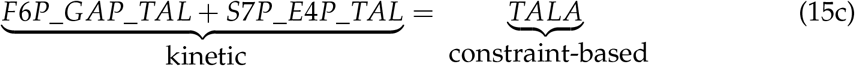

These equivalences are obtained by taking into account the cumulative effect of reactions. For instance, Equation (15b) is obtained by:

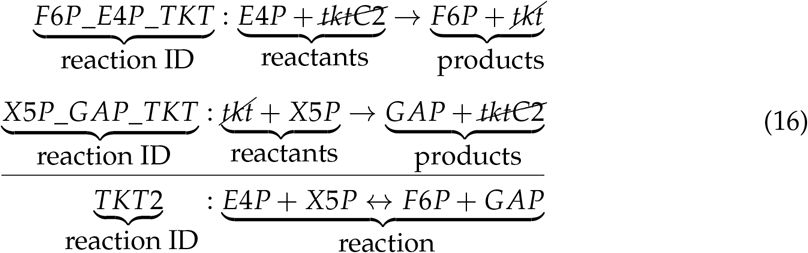

The sum of two reactions in the kinetic model, (*F6P_E4P_TKT* and *X5P_GAP_TKT*) results in reaction *TKT2* from the constraint-based model. This means that the kinetic model uses two reactions to produce metabolites *F6P* and *GAP* when *E4P* and *X5P* react. Two reaction intermediates are also specified (*tkt* and *tktC2*). In contrast, the constraint-based model ignores these intermediates and uses only one reaction to catalyze the same process. Similar differences also apply for the Equations (15a) and (15c).

To determine flux bounds, consider that under the steady state assumption, metabolite concentrations are assumed to be constant, i.e. the incoming flux to a metabolite must be equal to its outgoing flux. This assumption allows us to establish relationships among the steady-state fluxes of reactions in the network depicted in Figure **22**. In particular, the relationship in (17) can be deduced by assuming that the concentration of *tktC2* and *tkt* is constant:

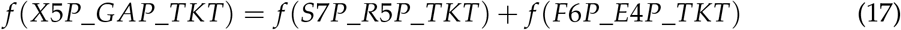

Similarly, equation (18) is obtained by considering metabolites *tal* and *talC3*:

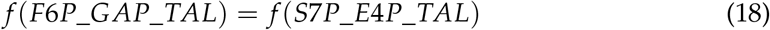

And, equation (19) is deduced from metabolite *S7P*:

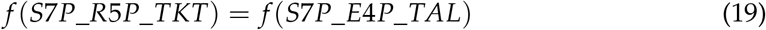

Metabolite *S7P* is only produced by reaction *S7P_R5P_TKT* in the kinetic model, and it is only produced by reaction *TKT1* in the constraint-based model. Thus, in order to further constrain the constraint-based model, the flux of *S7P_R5P_TKT* (kinetic model) can be used as follows to bound the flux *TKT1* (constraint-based model):

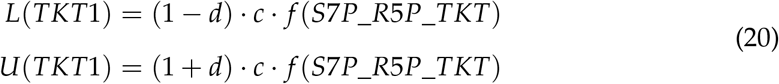

#### On the sequence of implemented kinetic bounds

In the main text, kinetic bounds were implemented according to Table **2**. To check the effect of the order in which bounds are implemented, Figure **24** shows the number of dormant reactions and the citramalate production flux under three different orders (the level of uncertainty is 10%). In particular, the implementation order for the results reported in Figure **24** is:

- Figure **24**(a): *SUCOAS, PGI, GND, FUM, SUCDi, PDH, G6PDH2r, PYK, TKT1, PFK, PGK, ICDHyr, CS, FBA, PGL, PGM, RPE, AKGDH, RPI, ENO, PPS, ACONTa, PPC, PPCK, FBP, GAPD, ME1, ACONTb, TPI*.
- Figure **24**(b): *PPS, SUCDi, FUM, RPI, CS, GAPD, G6PDH2r, ENO, PPC, RPE, PGM, SUCOAS, PGK, TPI, PDH, TKT1, GND, PYK, PPCK, ME1, ACONTb, PGI, PGL, ACONTa, FBA, FBP, ICDHyr, PFK, AKGDH*.
- Figure **24**(c): *PPCK, SUCOAS, PGL, GND, TPI, PYK, RPI, ENO, FBA, AKGDH, PGM, GAPD, PGI, FBP, ICDHyr, G6PDH2r, ACONTb, PPC, SUCDi, FUM, PPS, PFK, ME1, RPE, PGK, PDH, CS, TKT1, ACONTa*.

Note that the effect of permuting the order of implementation of the kinetic bounds in the citramalate-producing iML1515 model neither affected the final citramalate flux nor the number of dormant reactions. However, it did alter the specific point at which a significant activation of previously dormant reactions and a notable reallocation of fluxes occurred.

**FIG 24.**
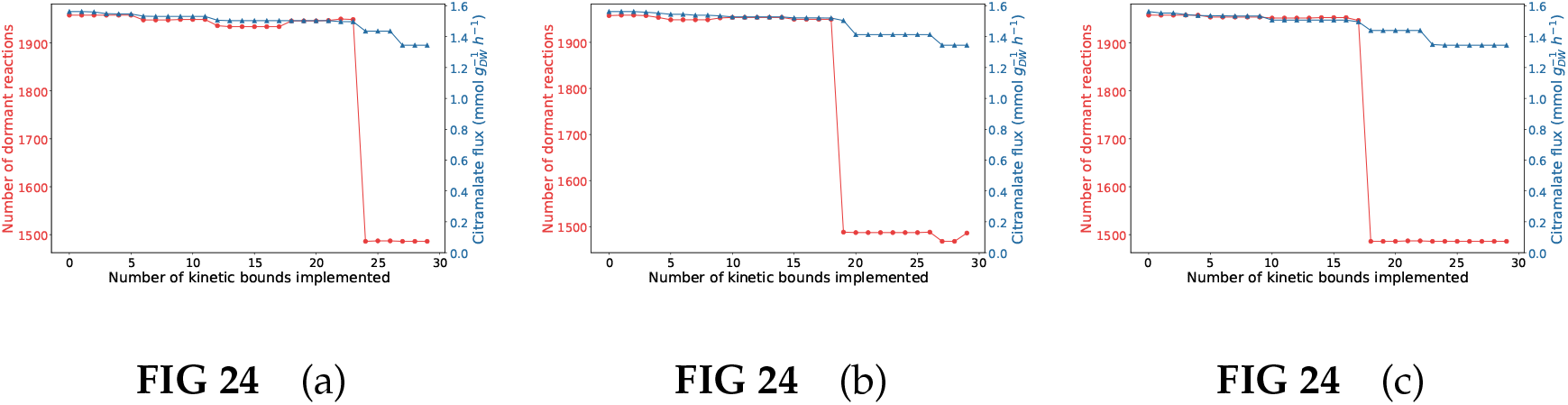
Dormant reactions (red line) and citramalate production flux (blue line) under three different orders.

### Additional file 1 — Mapping_Kinetic_Stoc.xlsx

This spreadsheet contains the mapping of the reactions between the genome-scale model and the reactions present in the kinetic model (“Mapping” sheet). “flux bounds” sheet contains the upper and lower bounds for each reaction in the genome-scale model that have equivalents in the kinetic model. “kinetic_fluxes_original” reports the fluxes of the reactions in the kinetic model after performing a simulation to reach the steady state. The units of the fluxes are expressed in *mM s*^−1^. Finally, “kinetic_fluxes_citra” reports the kinetic fluxes after a simulation of the citramalate-producing kinetic model until the steady state is reached. The units of the fluxes are expressed in *mM s*^−1^.

